# Combining Bulk and Single-Cell RNA Sequencing Data to Identify RNA methylation and Autophagy-Related Signatures in Patients with Chronic Obstructive Pulmonary Disease

**DOI:** 10.1101/2023.09.15.557860

**Authors:** Shixia Liao, Lanying Zhang, Yanwen Wang, Shuangfei He, Maomao Liu, Dongmei Wen, Jian Zhou, Yuting Liu, Pengpeng Sun, Qi Wang, Yang Xu, Yao OuYang

**Affiliations:** Department of Respiratory Medicine, Affiliated Hospital of Zunyi Medical University, Guizhou Province, 563003, China; West China Clinical Medical College, Sichuan University,Chengdu 610041, China; Department of Osteopathy, Affiliated Hospital of Zunyi Medical University, Guizhou, 563003, China; China-Canada Medical and Health Science Association, Toronto, L3R 1A3, Canada; Department of Anesthesiology, West China Hospital, Sichuan university, Chengdu 610041, China

**Keywords:** COPD, RNA methylation, Autophagy, bulk RNA sequencing, single cell RNA sequencing

## Abstract

**Background:** Chronic Obstructive Pulmonary Disease (COPD) is a heterogeneous lung condition associated with RNA methylation and autophagy. However, the specific autophagy-related genes and RNA methylation regulators involved in COPD development remain unknown.

**Methods:** We analyzed COPD and non-COPD patients datasets obtained from the Gene Expression Omnibus database, including Tissue Sequencing Transcriptome (bulk-seq) and single-cell sequencing (scRNA-seq) data. Differentially expressed genes (DEGs) were identified through differential genetic analysis using non-COPD bulk-seq data as the control group and COPD samples were used as the experimental group. Animal experiments were conducted to validate the expression of key genes. COPD model mice were exposed to smoke for four months, and lung function and histopathological changes were assessed. The mRNA and protein expression levels of *FTO, IGF2BP2, DDIT3, DNAJB1*, and *YTHDF3* were measured using RT-qPCR and Western blotting, respectively.

**Results:** We identified *FTO, IGF2BP2*, and *YTHDF3* as key methylation genes, along with autophagy hub genes *DDIT3* and *DNAJB1*. Animal experiments showed significantly increased mRNA and protein levels of *FTO, YTHDF3* and DNAJB1 and significantly decreased levels of *IGF2BP2* in lung tissue of COPD mice compared to the control group.

**Conclusion:** Our findings suggest that *DDIT3* and *DNAJB1* as autophagy hub genes, along with *FTO, IGF2BP2* and *YTHDF3* as RNA methylation genes, may play crucial roles in the development of COPD. These findings, supported by bulk-seq and scRNA-seq data, contribute novel genetic evidence for understanding the epigenetics of COPD.

## 1. Introduction

Chronic Obstructive Pulmonary Disease (COPD) is characterized by persistent, frequently progressive, airflow restriction that is caused by abnormalities of the airways and/or alveoli. More than 3 million individuals died from COPD in 2012, constituting 6% of total deaths worldwide and making it one of the top three causes of death (1, 2). Bronchodilator medicines have been the foundation of COPD treatment, with the additional option of inhaled corticosteroids to reduce COPD exacerbation. However, these drugs do not control COPD progression in all patients. In particular, this difficulty can be attributed to the understanding that COPD is a complex heterogeneous disease related to genetic changes (3). Therefore, its pathogenesis needs further study.

As a particularly important epigenetic modification, RNA methylation plays a key role in the regulation of tissue-specific gene expression (4, 5). Recent evidence has demonstrated that the expression of tumor-related genes is controlled by RNA regulators, which have a considerable impact on the formation and progression of cancers (6, 7). However, the roles of RNA regulators in COPD are yet to be explored.

Additionally, the breakdown of cytoplasmic proteins or lysosomal organelles is required for autophagy, a process of cell self-renewal. Several lung diseases are linked to abnormal or impeded autophagy (8). Chen et al. determined the presence of elevated levels of established autophagy markers (LC3-B, Atg4, and Atg7) in the lung tissue of COPD patients. Additionally, cigarette smoke extract (CSE) exposure has been observed to decrease histone deacetylase activity, thereby leading to an increase in the binding of the LC3B promoter to the early growth response-1 and E2F factors; contrastingly, epithelial cells are shielded from CSE-induced apoptosis by LC3B knockdown (9). Defects in autophagy lead to failure in the elimination of oxidative stress–damaged proteins and organelles, ultimately increasing the progression of COPD (10). Recently, there has been increasing evidence that COPD patients have dysregulated autophagy. Nevertheless, key autophagy genes and their involvement in the pathogenesis of COPD have rarely been reported. Therefore, studies focusing on key RNA modification and autophagy genes have contributed to deepening the understanding of COPD and will, consequently, have an important impact on the epidemiological control of this disease. The lack of reported genes is also a key problem in establishing a predictive model for the early diagnosis of COPD patients to improve quality of life and long-term survival rates.

Our study aimed to explore the changes in critical RNA methylation and autophagy regulators during COPD pathogenesis. We analyzed transcriptome data from COPD and non-COPD samples, as well as single-cell datasets, to identify differentially expressed genes enriched in the autophagy pathway and assess the expression of RNA methylation regulators. Furthermore, we identified key genes associated with autophagy and RNA modification and validated their expression levels in a COPD mouse model. This study provides insights into the role of RNA methylation and autophagy in COPD and may contribute to advancements in COPD diagnosis and treatment.

## Materials and Methods

### 2. Data collection

We utilized the COPD bulk-seq dataset GSE76705 from the official GEO website (https://www.ncbi.nlm.nih.gov/geo/) (11, 12) The GSE76705 dataset included 135 COPD samples and 229 non-COPD samples obtained from the ECLIPSE study (13-16). Additionally, we utilized the single-cell RNA sequencing datasets GSE167295(17) and GSE128033(18) from the GEO database (17, 18). The GSE167295 dataset included three COPD samples, and the GSE128033 dataset included three normal lung tissue samples (Figure 1).

**Figure 1.**
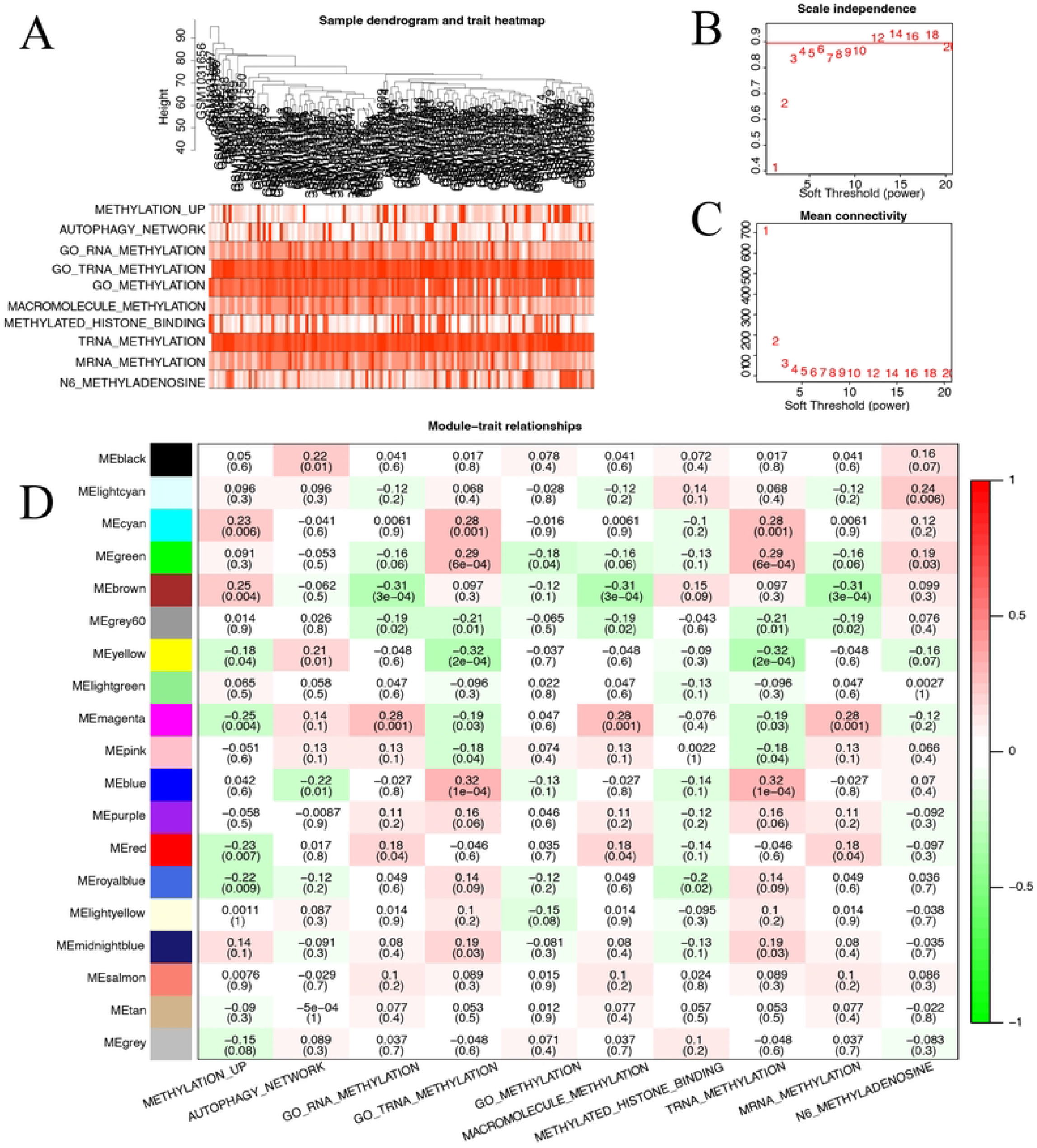
Flow chart of analysis conducted in this study. GSVA, Gene Set Variation Analysis; WGCNA, weighted gene co-expression network analysis; scRNA, single-cell RNA; DEGs, differentially expressed genes.

#### 2.1 Differential expression analysis of RNA-modified genes

We used the R package limma (version 3.52.2) for differential gene analysis of COPD and non-COPD samples from the GSE76705 dataset. To identify differentially expressed genes, the screening thresholds used for significantly differentially expressed genes was a p value < 0.05, and logFC > 1. By reviewing a previous study (19), 35 methylation-related RNA modifier genes were obtained, including 26 m6A regulators (*METTL3, RBM15B, ZC3H13, METTL14, METTL16, WTAP, RBM1, YTHDC1, YTHDC2, RBM15, YTHDF2, YTHDF3, KIAA1429, VIRMA, IGF2BP1, IGF2BP2, HNRNPG, LRPPRC, FTO, ALKBH5, HNRNPA2B1, HNRNPC, YTHDF1, IGF2BP3, HRBMX*, and *FMR1*) and 9m1A regulators (*TRMT10C, BMT2, RRP8, TRMT6, TRMT61A, TRMT61B, TRMT61C, ALKBH1*, and *ALKBH3*). Volcano and box plots were produced using the R package ggplot2(11); heat maps were plotted using the R package heatmap(12).

#### 2.2 Univariate logistic regression with the least absolute shrinkage and selection operator (Lasso) regression

We used univariate logistic regression to assess the association of 35 RNA-modified genes with COPD in the GSE76705 dataset; the expression data were processed using “scale” for the expression matrix of RNA-modified genes, followed by univariate logistic regression with the presence or absence of COPD. Significant genes, with p-values < 0.05, were selected for further lasso regression.

#### 2.3 Gene localization map

For the GSE76705 dataset, the GENCODE database(20), based on the human reference genome function “UCSC.HG19.Human.CytoBandIdeogram”, was used to query the location of genes that were associated with COPD and passed Lasso regression on chromosomes; then, we used the R package RCircos(21) (version 1.2.2) to map the chromosomal locations of these genes obtained by Lasso regression.

#### 2.4 Gene Ontology (GO) enrichment analysis

GO enrichment analysis (22) of RNA-modified genes in the GSE76705 dataset was performed by using the R package clusterProfiler (23). For each GO entry, the differential gene count was tallied, and a hypergeometric distribution technique was used to determine the importance of the differential gene enrichment in each entry. The output from the corresponding calculation was a p-value according to the significance of the enrichment; a p-value < 0.05 was considered statistically significant. Bubble and network plots were produced for the top 10 enrichment results.

#### 2.5 GSEA

We utilized the Molecular Signatures Database (MSigDB) gene set, msigdb.v7.0.entrez.gmt, for GSEA analysis (24). The R package clusterProfiler (version 4.4.4) was used with threshold values, including organization = “hsa”, pvalueCutoff = 0.2, nPerm = 1000, minGSSize = 10, maxGSSize = 500, verbose = TRUE, seed = FALSE, and by = “fgsea” (23). The top 15 pathways, including autophagy-related pathways, were selected for mountain range mapping and gene enrichment mapping.

#### 2.6 Gene set variation analysis (GSVA)

We utilized GSVA (25) with the “gsva” method to analyze single-cell data from GSE167295 and GSE128033. Gene sets from the MSigDB database (24) were employed. GSVA adjusted the gene expression data, converting single-gene features to gene-set features. Enrichment Scores were calculated for each gene set, representing pathway enrichment in each cell. Limma (26) identified pathways with significant differences (p-value < 0.05 and logFC > 1), and pathway activity scores were compared between cell groups. A heat map was generated using the top 20 pathways ranked by t-value.

#### 2.7 WGCNA

WGCNA, is a data mining technique suitable for biological network research based on correlations between variables(27). We applied WGCNA to the GSE76705 dataset, selecting the top 25% of genes with the highest expression variance. Using the adjacency matrix and topological overlap matrix, we performed hierarchical clustering to identify gene modules with a minimum module membership of 30. The GSVA enriched pathways were integrated with each module to examine their functions and generate a module function heat map. We identified key genes for RNA modification by intersecting the modules associated with methylation modification and the previously obtained RNA modification genes.

#### 2.8 CIBERSORT immuno-infiltration analysis

CIBERSORT is an algorithm that utilizes linear support vector regression to estimate the composition and abundance of immune cells in a transcriptome expression matrix(28). We applied CIBERSORT to the GSE76705 dataset, transforming the data into absolute abundance of immune cells and stromal cells. This allowed us to evaluate 22 types of immune cell subpopulations. The samples were filtered based on a p-value threshold of <0.05 to generate an immune cell infiltration matrix. Subsequently, data with immune cell enrichment fractions >0 were further filtered to obtain the final results of specific immune cell infiltration. We used the R package pheatmap for plotting the correlation heat map and the R package ggpubr for plotting the correlation scatter plot (29).

#### 2.9 Quality control of COPD single-cell data using Seurat

First, we used R (https://www.r-project.org/, version 4.2.1) with the R package Seurat (version 4.1.1) (30). We created Seurat objects from the merged expression matrix of the GSE167295 and GSE128033 datasets. It must be noted that the proportion of mitochondrial genes in all genetic material may indicate whether a cell is in homeostasis. Generally, we assumed that a cell would be under stress if it contained a higher proportion of mitochondrial genes than all other genes; thus, we eliminated cells with > 5 mitochondrial genes. Since duplicate cells may have an excessively high number of genes and low-quality cells or empty droplets typically have few genes, we selected cells with the corresponding parameters: nFeature_RNA > 200, nCount_RNA < 3000, and percent. mt < 5.

#### 2.10 SingleR cell type annotation

To identify differential genes between cell types, we utilized cell type annotation of single-cell data using the BlueprintEncodeData (31) dataset from the R package SingleR (version 1.10.0)(32) across the following cell types: T-cells, B-cells, NK cells, plasma cells, epithelial cells, and myeloid cells (DC, macrophages, monocytes, stromal cells, mast cells, and endothelial cells). We selected CD8 T-cell-related genes and B-cell-related genes and observed the expression of these gene sets between clusters using clustering and violin plots.

#### 2.11 Differential expression of RNA-modified genes between cells

Next, we used the “FindAllMarkers” function, which compares the differences in gene expression between clusters using a Wilcoxon rank-sum test. We intersected the RNA-modified genes with the cluster marker genes to determine the RNA-modified genes that were differentially expressed among these clusters. The heat map was plotted using the “DoHeatmap” function, the violin map was produced using the “FeaturePlot” function, and the mountain map was plotted using “RidgePlot”. This analysis was conducted to establish the expression of these RNA-modified genes in different cells.

#### 2.12 Intercellular differential expression of autophagy genes

We extracted 232 genes involved in autophagy from the Human Autophagy Database (33-35) (http://www.autophagy.lu/) and intersected them with cell cluster marker genes to determine the differentially expressed genes between cell clusters. The autophagy-related genes were extracted and intersected with the cluster marker genes to obtain the autophagy-related genes that were differentially expressed among these cell clusters. To show the expression of these autophagy-associated genes in different cells, the corresponding heat map was plotted using “DoHeatmap” function, the cluster map was produced using the “FeaturePlot” function, and the mountain map was plotted using “RidgePlot”.

#### 2.13 Cell Chat

CellChat is a tool for examining and contrasting intercellular communication networks. Using scRNA-seq data, CellChat analyzes the expression of signaling ligands, receptors, antagonists, soluble agonists, and inhibitory or stimulatory membrane-bound co-receptors to identify intercellular communication networks (36). To identify these cell–cell communication networks, we utilized a list of human receptor–ligand pairs from the R package database ‘CellChatDB.human’ (37). This information was then integrated with gene expression information, with lines connecting two cells sharing a receptor– ligand pair; finally, this output was used to generate a cell communication map.

#### 2.14 Real-time quantitative polymerase chain reaction (RT-qPCR)

Messenger (m) RNA expression of FTO, IGF2BP2, DDIT3 and DNAJB1 were evaluated by RT-qPCR using an ABI7500 instrument (Applied Biosystems, Foster City, United States). Total RNA was extracted from mice lung tissue (100 mg) with TRIzol Reagent. The purity and concentration of RNA were determined. Complementary (c)DNA synthesis was done using a cDNA synthesis kit (Thermo, USA). The primer sequences were listed in Table S1. we used (forward and reverse, respectively) were: 5′-ATGTTGAAGATGAGCGGGTGG-3 ′and 3 ′-ATGTGCGTGTGACCTCTGT T-5 ′for DDIT3;5′-CATTCCGGGGATCTAGCGGC-3and 3′-CCGAAGGTCATGGGTGACTG -5′ for DNAJB1;5′-GAGCAGCCTACAACGTGACT-3′and 3′-TATACTGGTGAGACCGGCC A-5′for FTO;5′-GGAGAACGTGGAGCAAGTCA-3′and 3′-CGCAGCGGGAAATCAATC TG-5′for IGF2BP2 and 5′-CACGCCTACAGATCCCACAG-3′and 3′-TCTGAGCCTCGT CACCTACA-5′for β-actin. SYBR Premix Ex Taq™ II was used for RT-qPCR. Quantitative data are expressed as relative expression according to the formula2^-△△ct.^

#### 2.15 Physiological testing of pulmonary function and Immunohistochemistry

Male C57Bl/6 mice (n = 12), aged three months, were divided into two groups: a control group and a smoked group. The mice were obtained from the Laboratory Animal Center at Third Military Medical University and were housed in ventilated polyethylene cages with controlled light, temperature, and humidity conditions. To induce COPD, the smoked group was exposed to smoke from 15 cigarettes, five days a week, for a duration of four months. The smoke was delivered into a plastic box where the mice were kept, allowing for passive inhalation. Prior to euthanasia, the mice were anesthetized with ketamine (100 mg/kg) and xylazine (10 mg/kg) administered intraperitoneally. All animal experiments were conducted in accordance with the ethical guidelines of Zunyi Medical University, approved by the Animal Ethics Committee (approval number: ZMU20202409). The experimental procedures followed the Guide for Care and Use of Laboratory Animals (NIH publication, revised 1996). HE staining and immunohistochemistry were performed to examine the localization and expression of selected genes in mice lung tissues.

#### 2.16 Statistical analysis

All data calculations and statistical analyses were performed using R software (https://www.r-project.org/, version 4.2.1). For the comparison of two groups with continuous variables, statistical significance of normally distributed variables was estimated using an independent Student’s t-test; differences between non-normally distributed variables were analyzed using a Mann-Whitney U-test (i.e., Wilcoxon rank sum test). A chi-square test or Fisher’s exact test was used to compare and analyze the statistical significance between the two sets of categorical variables. Finally, correlation coefficients were calculated between different genes using Pearson correlation analysis.

## 3. Results

### 3.1 Expression profiling of RNA-modified genes

In this study, we analyzed 364 samples from the GSE76705 dataset, including 229 non-COPD samples and 135 COPD samples. Differential gene analysis was performed using the COPD samples as the experimental group and the non-COPD samples as the control group. We identified 28 RNA modification genes based on a previous study and obtained 14 differentially expressed genes in COPD tissues (threshold p < 0.05, logFC ≥ 0.5). Among the upregulated genes were KIAA1429, IGF2BP2, LRPPRC, TRMT61A, HNRNPC, and RRP8, while the downregulated genes included TRMT61B, YTHDC1, TRMT10C, ZC3H13, METTL3, FMR1, YTHDF3, and METTL14. Protein-protein interaction analysis revealed associations of these genes with various RNA modification proteins. A volcano plot and correlation heat map were generated to visualize the differential expression and correlation patterns of these genes. The expression profiles of the RNA-modified genes were presented in a heat map, and the expression levels of the differentially expressed genes were depicted in a bar graph. The results highlight the dysregulation of RNA modification genes in COPD. (Figure 2A-E)

**Figure 2.**
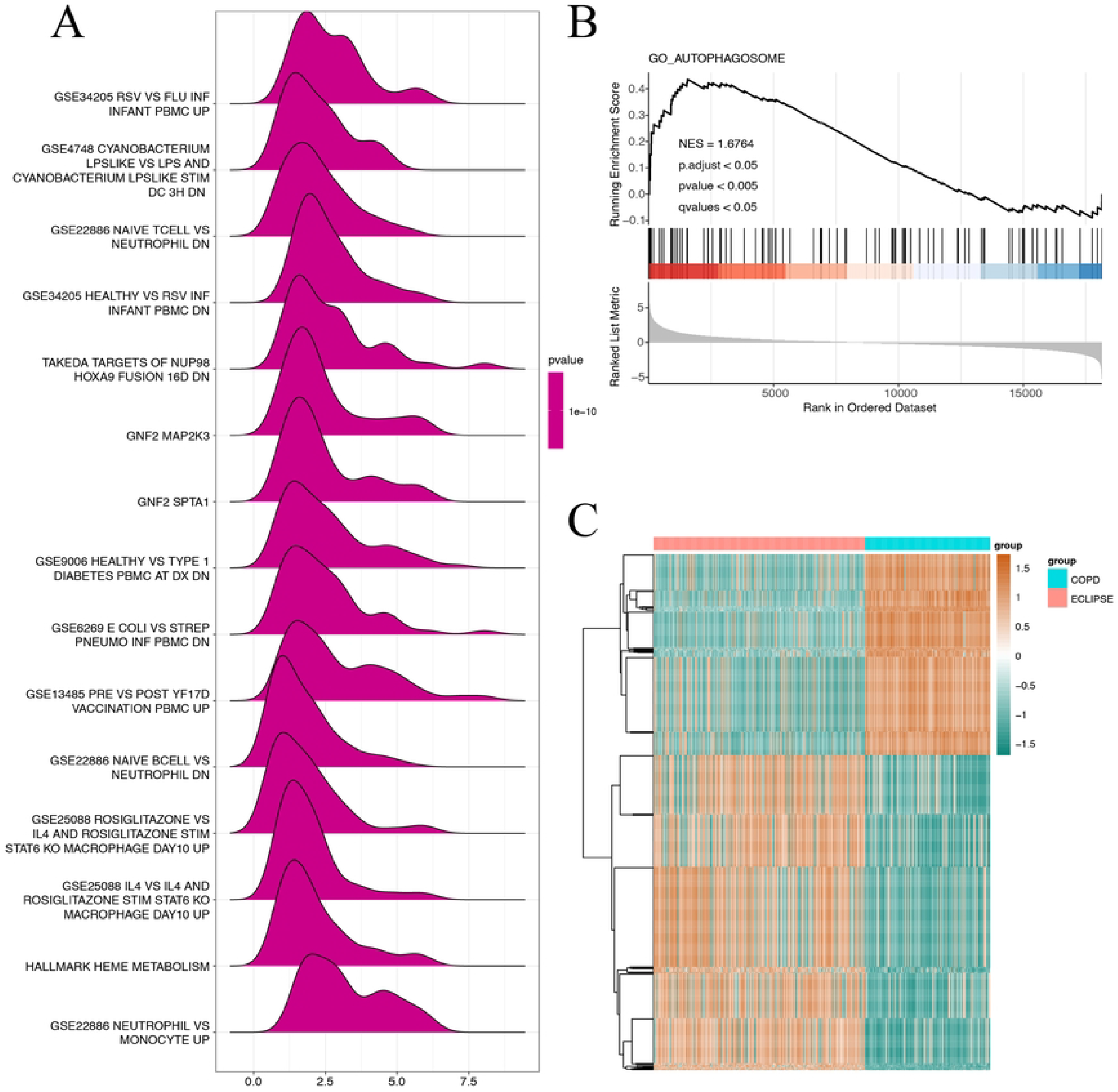
RNA modifier gene expression in COPD and non-COPD groups. (A) Network mapping of 15 selected RNA-modified genes in Cytoscape. (B) Volcano plot of 14 differentially expressed genes. (C) Heat map of 28 differentially expressed genes. (D) Correlation heat map of 14 genes. (E) Bar graph showing differential expression. *p < 0.05, **p < 0.01, ***p < 0.001, ****p < 0.0001. COPD, chronic obstructive pulmonary disease. Red denotes up-regulation and blue, green, and yellow denotes down-regulation.

### 3.2 RNA-modified genes in the disease group/control group distinction

To investigate the association of RNA modifier genes with COPD in the GSE76705 dataset, logistic regression analysis was performed on 28 RNA modifier genes, resulting in 25 genes passing the univariate logistic regression analysis. Lasso regression analysis was then conducted on these 25 genes, which identified 13 significant genes (METTL14, KIAA1429, ZC3H13, FTO, YTHDC1, YTHDF2, YTHDF3, IGF2BP2, IGF2BP3, HNRNPC, TRMT10C, RRP8, and ALKBH1) (Figure 3A-B). The chromosomal locations of these 13 genes were determined using the GENCODE database (version v22) (20), and a gene chromosome localization loop (Figure 3C) was generated using the RCircos R package (version 1.2.2)(21). The corresponding chromosomal loci were as follows: YTHDF2 on chromosome 1, TRMT10C and IGF2BP2 on chromosome 2, YTHDC1 and METTL14 on chromosome 4, IGF2BP3 on chromosome 7, YTHDF3 and KIAA1429 on chromosome 8, RRP8 on chromosome 11, ZC3H13 on chromosome 13, ALKBH1 and HNRNPC on chromosome 14, and FTO on chromosome 16.

**Figure 3.**
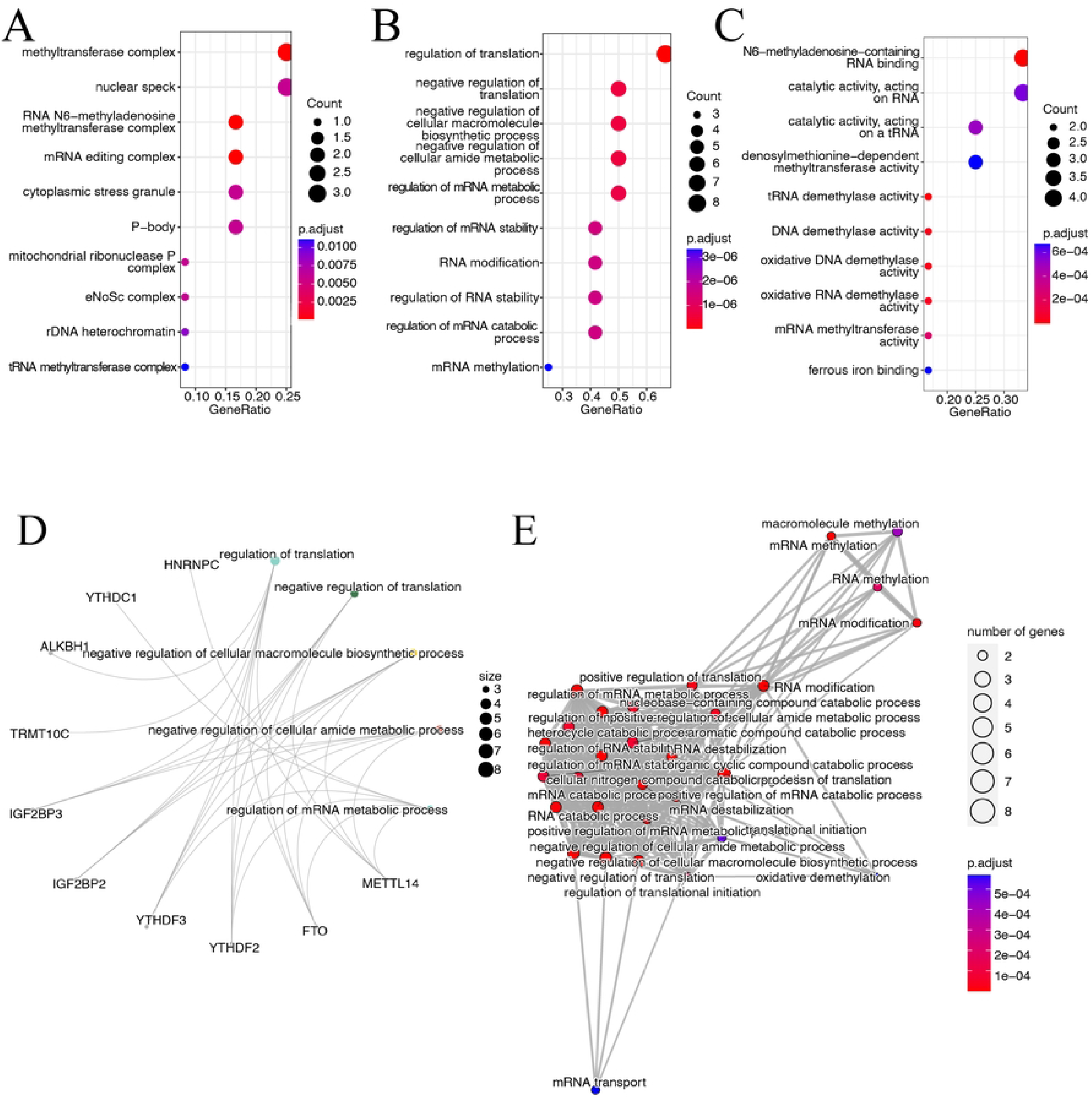
Logistic regression, LASSO regression, and gene localization plots for 13 RNA-modified genes in the GSE76705 dataset. (**A**) Regression coefficient plots showing the sequential decreasing coefficients of each variable as the penalty coefficient λ increases. (**B**) Point box plot showing that the trend of the overall model error varies as the penalty coefficient λ increases. (**C**) The loci of the 13 RNA-modified genes on the chromosomes. LASSO, least absolute shrinkage and selection operator.

### 3.3 Enrichment analysis of RNA-modified genes

GO enrichment analysis of the 13 RNA modification genes in the GSE76705 dataset revealed important insights. Figure 4A shows the regulation of translation and macromolecular biosynthetic processes, while Figure 4B highlights the regulation of translation in biological processes. These findings indicate significant upregulation of mRNA methylation. In terms of molecular function, N6-methyladenosine-containing RNA binding and various methyltransferase activities were observed (Figure 4C). The relationship between enriched signaling pathways and RNA modification genes is depicted in Figure S1A. Notably, certain genes were downregulated in specific processes, such as HNRNPC in mRNA catabolism and IGF2BP2 in regulating translation (Figure 4D). The connection between enriched pathways, particularly those related to mRNA methylation, is illustrated in Figure 4E.

**Figure 4.**
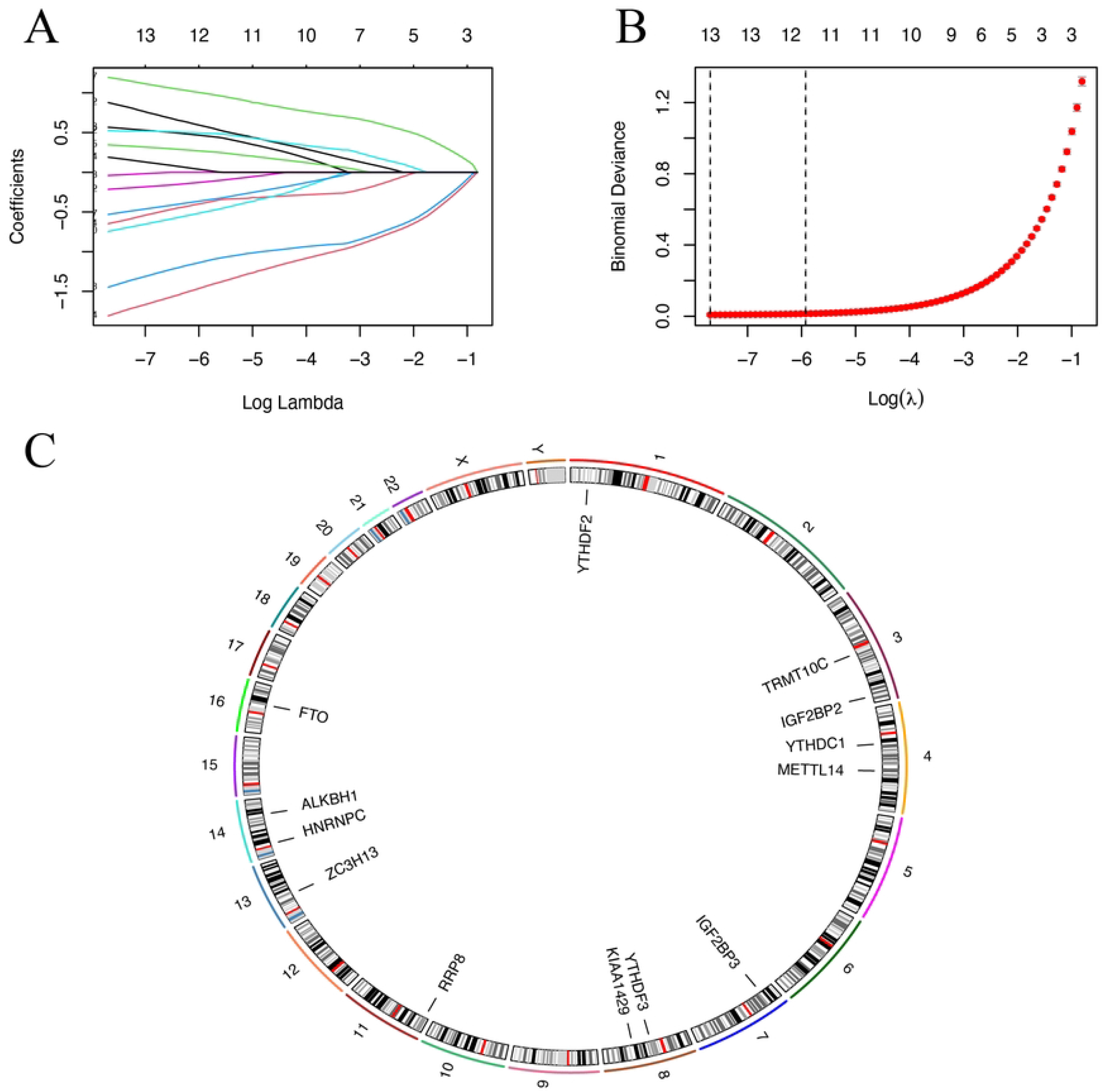
Enrichment analysis of RNA-modified genes in the GSE76705 dataset. (**A-C**) The gene ontology (GO) enrichment bubble plots for the 13 RNA modified genes. (**D**) Network diagram of top 5 signaling pathway and related genes in the enrichment pathway; the size of the pathway nodes represents the number of links to each gene. (**E**) Protein–protein network interaction map; the size of the nodes represents the number of enriched genes. The color represents the p-value: the bluer the color the greater the p-value, the redder the color the smaller the p-value.

### 3.4 GSEA and GSVA enrichment analysis

In the GSE76705 dataset, differential gene analysis revealed enrichment of immune cell-related pathways and autophagy-related signaling pathways (Figure 5A-B). GSVA analysis of 13 RNA modification genes identified multiple differentially enriched pathways, including RNA methylation and mRNA modification (Figure 5C). Significantly enriched pathways included CAGCCTC MIR4855P, Schlosser serum response dn, GSE26669 CD4 vs. CD8 T-cell in MLR costim block up, and GSE45739 NRAS KO vs. WT ACD3 ACD28 STIM CD4 T-cell dn (log FC > 1.5 and p-value < 0.05).

**Figure 5.**
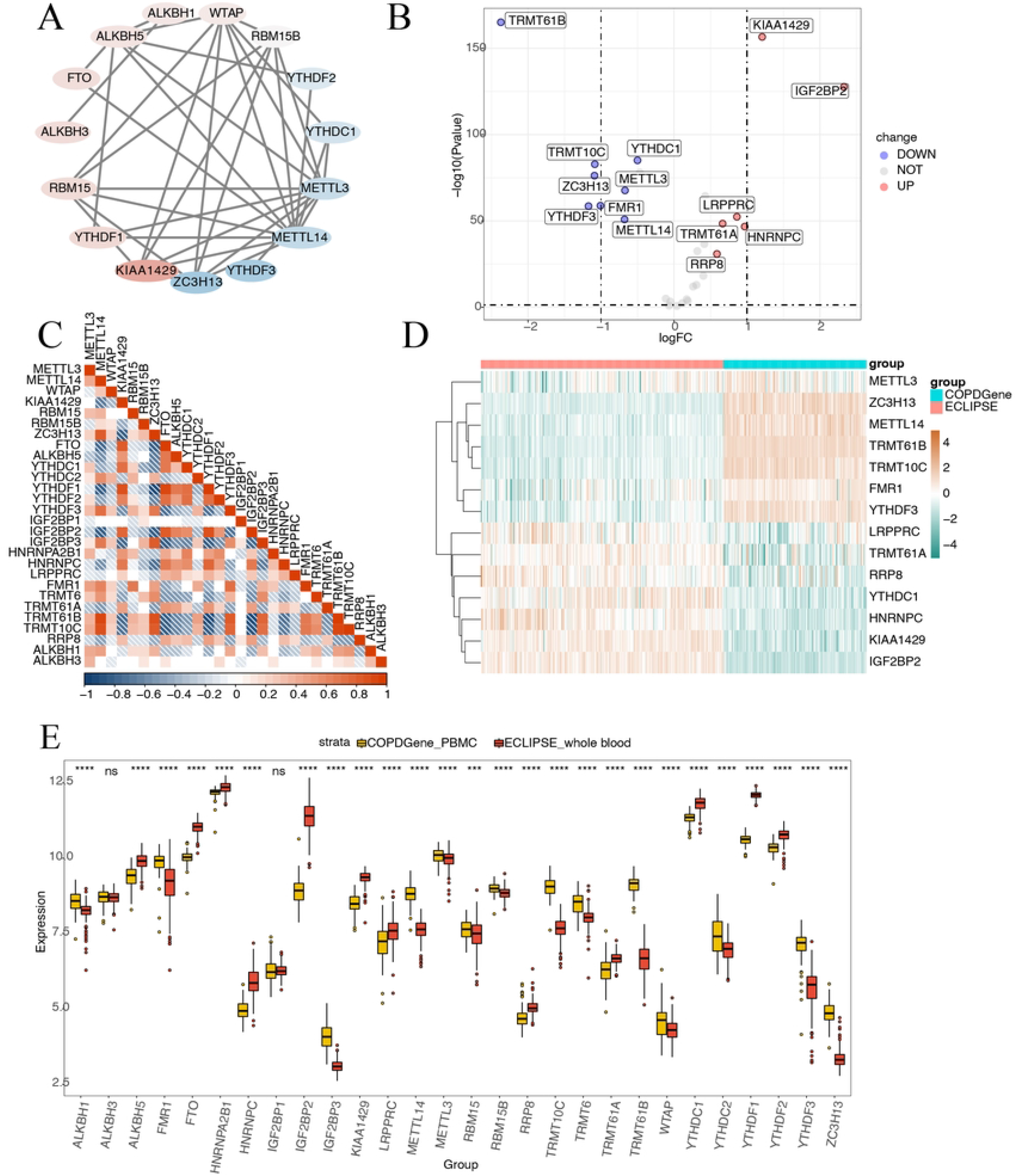
Gene set enrichment analysis (GSEA) and gene set variation analysis GSVA of the gene expression data from the GSE76705 dataset. (**A**) The top 15 GSEA enrichment results; the x-axis denotes the enrichment score, the y-axis denotes the name of the pathway, and the shade of the fill color denotes the size of the p-value. (**B**) GSEA enrichment graphs of autophagy-related pathways. NES: the maximum enrichment fraction, p value: the original p value, p.adjust: the corrected p value, and qvalues: the p value after correction with the false discovery rate method. (**C**) Heat map of differential enrichment from GSVA enrichment results in COPD and ECLIPSE groups. Red denotes up-regulated and green denotes down-regulated. COPD, chronic obstructive pulmonary disease.

### 3.5 WGCNA

Using WGCNA analysis, we constructed co-expression networks and identified highly related genes. The clustering heat map demonstrated sample clustering based on autophagy-related pathways and methylation-related enrichment pathways (Figure 6A). The soft threshold β value of 4 was chosen to construct a scale-free network (Figure 6B-C). Nineteen modules were identified, and the module correlation heat map was visualized (Figure 6D). From the green and blue modules, we identified the YTHDF3 gene by intersecting genes with methylation pathway correlation and RNA modification genes. From the black and yellow modules, we identified DNAJB9, NAMPT, DNAJB1, PPP1R15A, CDKN1A, CXCR4, DDIT3, VEGFA, FKBP1B, and NCKAP1 genes by intersecting genes with autophagy pathway correlation and autophagy marker genes.

**Figure 6.**
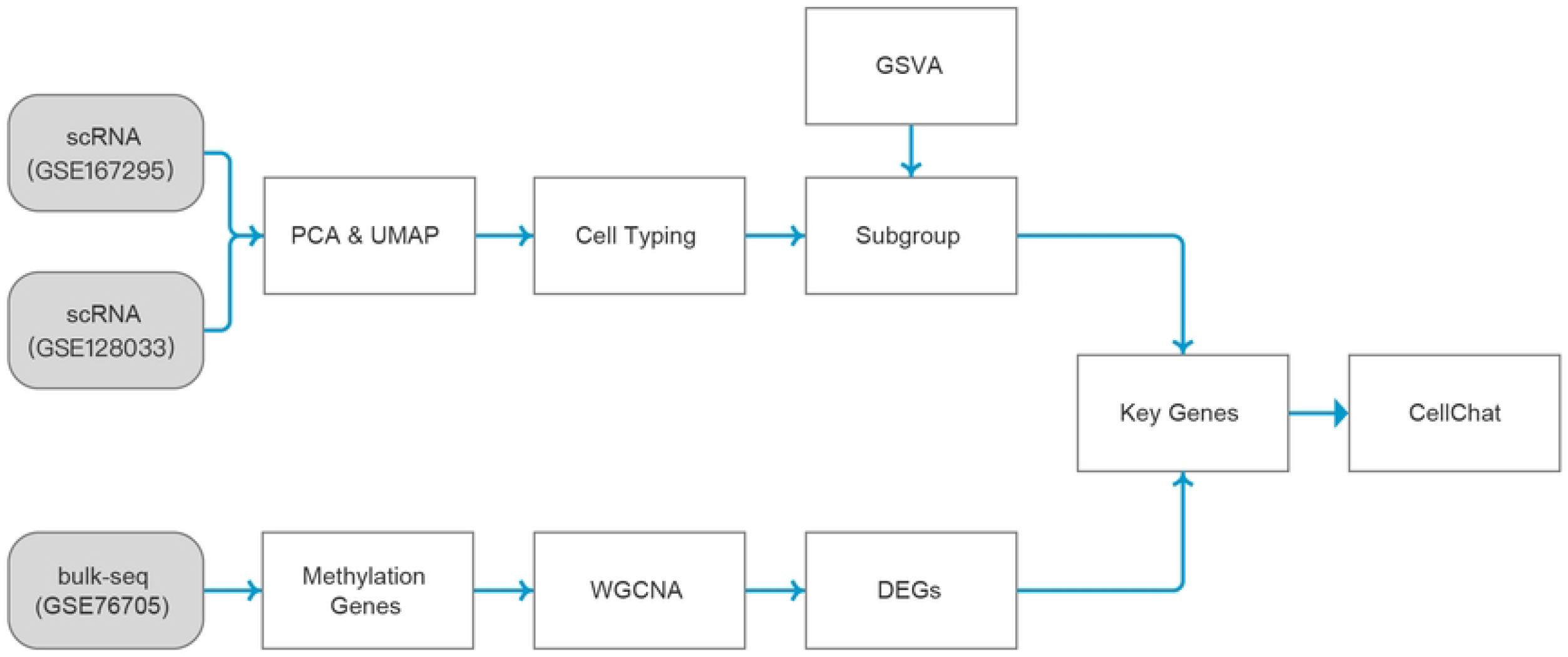
Weighted gene co-expression network analysis (WGCNA) modules associated with cell types in the GSE76705 dataset. (A) Sample clustering and enrichment heat map. (B) Scale-free fit index analysis. (C) Average connectivity analysis. The horizontal axis represents the weight parameter β and the vertical axis represents the mean value of all gene neighboring functions in the corresponding gene module. (D) Module correlation heat map with methylation and autophagy phenotypes. The correlation coefficients and p-values of the corresponding immune cells with the modules are provided in each box.

### 3.6 Immuno-infiltration analysis

We analyzed the correlation between the 13 RNA-modified genes and infiltrated immune cells using the CIBERSORT algorithm. Thirteen immune cell types were identified, including memory B-cells, naive B-cells, plasma cells, CD8 T-cells, CD4 memory activated T-cells, CD4 naive T-cells, γδT-cells, regulatory T-cells, monocytes, resting NK cells, resting mast cells, M2 macrophages, and neutrophils. The correlation heat map (Figure 7A) and scatter plots (Figure 7B-I) showed significant correlations between certain genes and immune cell types. For example, monocytes showed a positive correlation with ZC3H13 and a negative correlation with FTO, while neutrophils exhibited various correlations with METTL14, KIAA1429, ZC3H13, FTO, IGF2BP2, and IGF2BP3.

**Figure 7.**
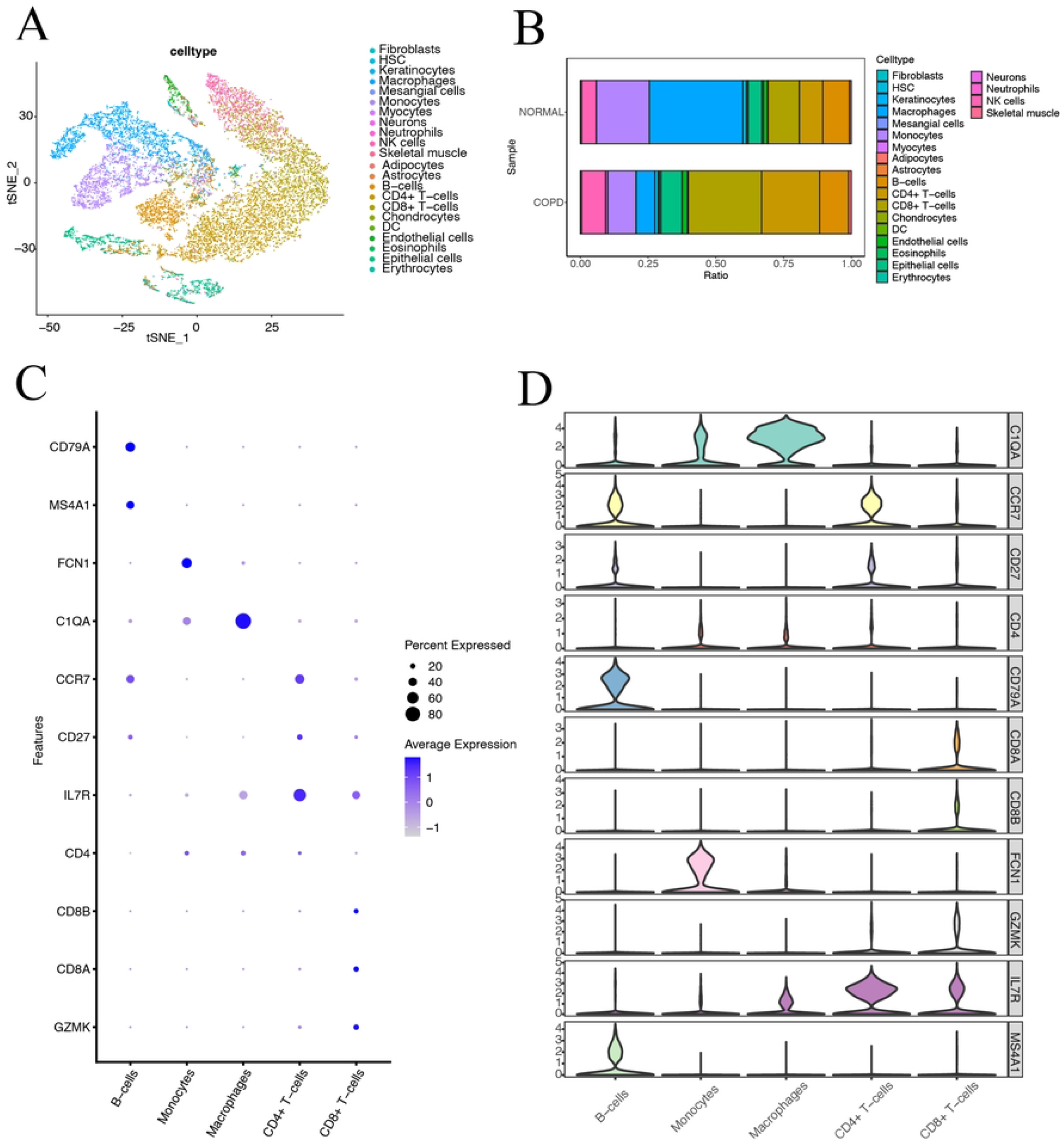
Correlation analysis of RNA-modified genes with infiltrating immune cells and stromal cells in the GSE76705 dataset. (A) Heat map of correlations between RNA-modified genes and immune cells/stromal cells. (B–I) Scatter plots showing correlations of RNA-modified genes with specific immune cell types. (B) Monocytes and ZC3H13 (r = 0.827, p < 0.001). (C) Monocytes and FTO (r = -0.858, p < 0.001). (D) Neutrophils and MET (r = -0.858, p < 0.001), Neutrophils and METTL14 (r = -0.815, p < 0.001). (E) Neutrophils and KIAA1429 (r = 0.831, p < 0.001). (F) Neutrophils and ZC3H13 (r = -0.875, p < 0.001). (G) Neutrophils and FTO (r = -0.875, p < 0.001), Neutrophils and FTO (r = 0.812, p < 0.001). (H) Neutrophils and IGF2BP2 (r = 0.884, p < 0.001). (I) Neutrophils and IGF2BP3 (r = -0.812, p < 0.001). Red indicates positive correlation, blue indicates negative correlation, and white indicates no significant correlation.

### 3.7 Single-cell data reveal cellular heterogeneity in COPD

Single-cell RNA data from COPD and non-COPD samples were analyzed, resulting in 18,523 cells after filtering. Violin plots (Figure 8A-C) illustrated gene expression, expression value, and mitochondrial gene expression proportion. PCA analysis (Figure 8D-E) selected 7 principal components, and tSNE clustering classified cells into 22 clusters. Tissue clustering maps (Figure 8F) showed the distribution of these clusters, and the clustering of COPD and normal lung tissue is shown in Figure 8G. Cluster 0 (17.534%) and cluster 1 (18.443%) were dominant in COPD cells, while cluster 5 (15.355%) and cluster 10 (10.626%) were predominant in normal cells.

**Figure 8.**
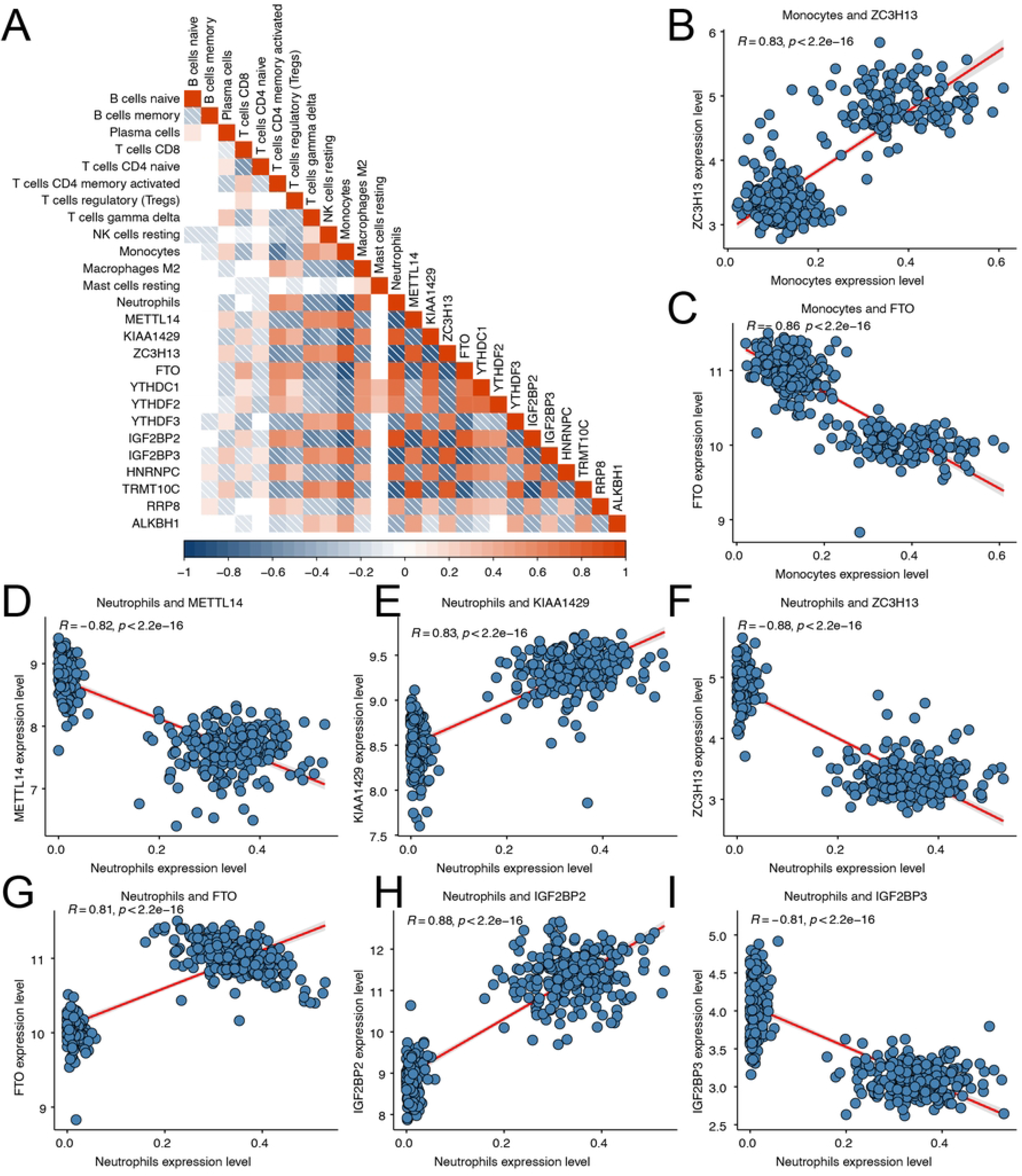
A total of 23 cell clusters with different annotations were identified using single-cell RNA-seq data, revealing the presence of high cellular heterogeneity in chronic obstructive pulmonary disease (COPD) sample cells. (**A–C**) Cells with nFeature_RNA > 200, nCount_RNA < 3,000, and per cent.mt < 5 were selected for quality control violin mapping. (**D**) The overall data distribution was observed using PC1 and PC2. (**E**) The relationship between the number of principal components and standard deviation. (**F**) Visualization of 22 clusters using the t-distributed stochastic neighbor embedding (tSNE) dimensionality reduction algorithm. (**G**) Visualization of COPD and normal tissue-derived cells using the tSNE dimensionality reduction algorithm.

### 3.8 Cell annotation and expression of cell population marker genes

We identified 22 cell types using SingleR, including adipocytes, astrocytes, CD4+ T-cells, CD8+ T-cells, B-cells, chondrocytes, DC, endothelial cells, eosinophils, epithelial cells, erythrocytes, fibroblasts, HSC, keratinocytes, macrophages, mesenchymal cells, monocytes, myocytes, neurons, neutrophils, NK cells, and skeletal muscle cells (Figure S1B). B-cells, CD4+ T-cells, CD8+ T-cells, macrophages, and monocytes showed significant upregulation, while T-cells and NK cells exhibited a negative correlation with macrophages and monocytes. The clusters of these cell types are illustrated in Figure 9A, with distinct annotations for each cluster. We also examined the distribution of cell types in COPD and normal tissues, highlighting the prevalence of specific cell populations in each (Figure 9B). The cell annotation was performed based on marker genes and previous studies: CD8+ T-cells (GZMK, CD8A, CD8B) (38), CD4+ T-cells (CD27, CCR7)(32), macrophages (C1QA) (38), monocytes (FCN1) (39), B-cells (MS4A1, CD79A) (40). Bubble and violin plots were generated to confirm the relatively high expression of marker genes in the corresponding cell types (Figure 9C-D), validating the accuracy of cell annotation. Bubble and violin plots were generated to confirm the relatively high expression of marker genes in the corresponding cell types (Figure 9C-D), validating the accuracy of cell annotation.

**Figure 9.**
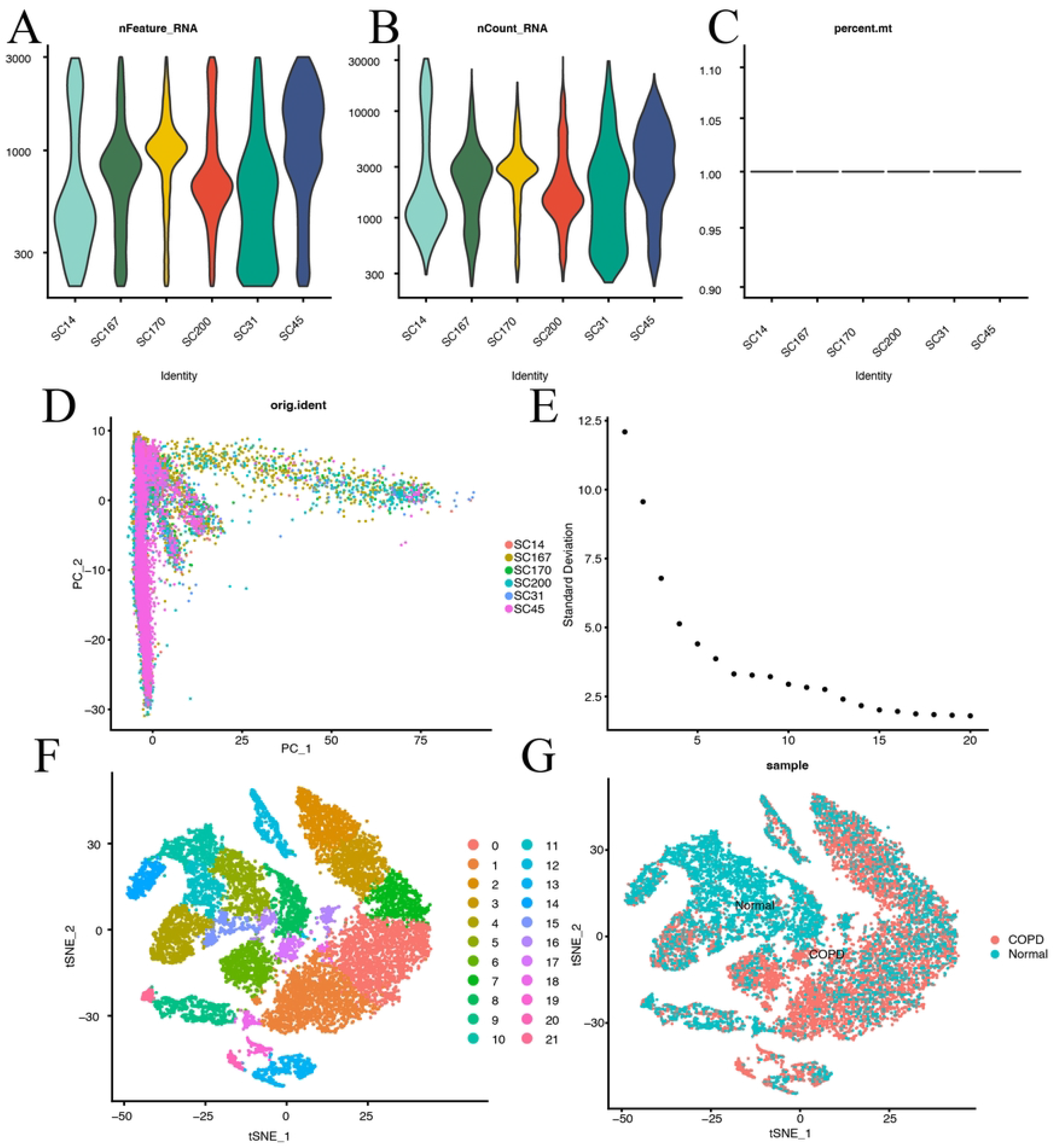
Cellular heterogeneity in junctional COPD cells. (A) tSNE cluster diagram of 22 cell clusters. (B) Cell type distribution in COPD and normal tissues. (C) Expression levels of specific genes across cell types shown in bubble plots. (D) Violin plot of gene expression levels across cell types.

### 3.9 Differential gene analysis between cell clusters

We re-clustered the five cell types (CD4+ T-cells, CD8+ T-cells, B-cells, macrophages, and monocytes) to form 21 independent clusters (Figure 10A). The distribution of cells within each cluster was analyzed, and the top differentially expressed genes in each cell type were identified (Figure 10B). The corresponding heat map was produced (Figure 10C). However, there was no intersection between the differentially expressed genes and RNA modification genes. Intercellular communication analysis revealed that macrophages and monocytes had stronger interactions with other cell types, particularly B-cells, CD4+ T-cells, and CD8+ T-cells (Figure 10D). The intensity of intercellular communication was visualized using heat maps and signal plots, indicating the contribution of specific pathways such as IL-1, IL-6, and resisting. We further analyzed the differential genes within the 21 clusters and identified autophagy-related and RNA modification-related intersection genes. Autophagy and methylation scoring were applied to macrophages, with A+ indicating high autophagy scores and M+ indicating high methylation scores. A+M+ macrophages exhibited increased frequency and intensity of interactions with other cell types compared to macrophages with low autophagy and methylation scores (Figure 10E and 10F).

**Figure 10.**
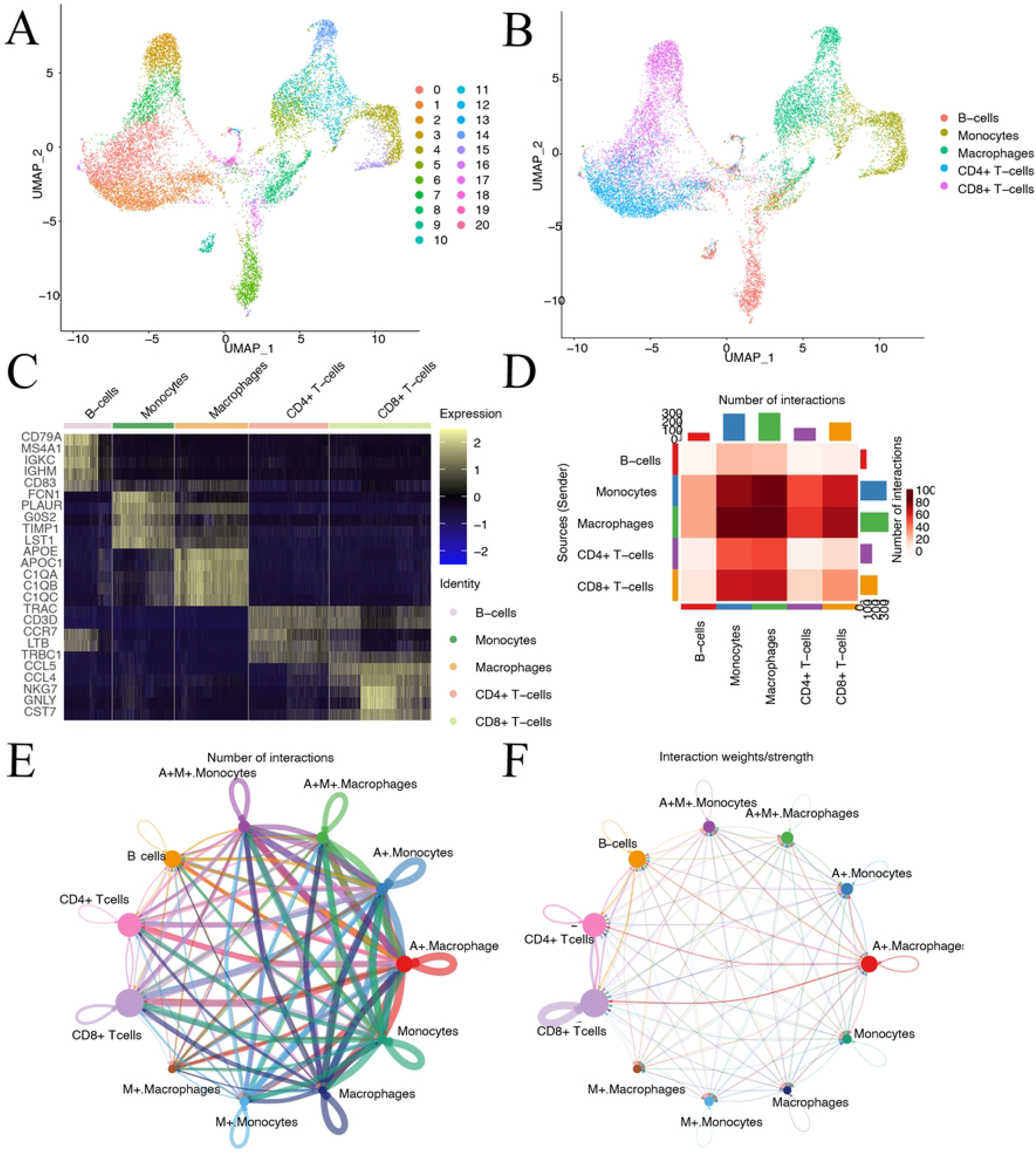
Expression profiles of 21 clustered differential genes with cellular communication analysis. (**A**) A total of 21 clusters containing B-cells, macrophages CD4+ T-cells, CD8+ T-cells, and monocytes. (**B**) Clustering distribution of five cell types in these 21 clusters. (**C**) Heat map of expression of the top 20 genes for differential gene analysis of the 21 clusters. (**D**) Heat map of the communication intensity of the five cell types. Macrophages and monocytes interacted with B-cells, CD4+ T-cells, and CD8+ T-cells with higher intensity. (**E**) Circle plot of cell communication data. (**F**) Round plot of cell communication intensity.

### 3.10 Experimental verification of the expression level of hub genes in mouse lung tissue

After 4 months of smoking, the model group exhibited a significantly lower FEV20/FVC(%) compared to the air group, indicating impaired lung function (Figure 11A). Histopathological analysis of lung tissue revealed normal alveolar structure and minimal inflammation in the control group, while the COPD group exhibited epithelial shedding, cilia damage, inflammatory cell infiltration, smooth muscle hyperplasia, and alveolar wall destruction (Figure 11B). mRNA expression levels of FTO, YTHDF3, and DNAJB1 were significantly up-regulated in COPD mice, while IGF2BP2 expression was significantly down-regulated, and DDIT3 expression showed no significant difference (Figure 11C). Protein expression levels of FTO, YTHDF3, and DNAJB1 were significantly increased, while IGF2BP2 expression was significantly decreased in COPD mice compared to control mice (Figure 11D-E).

**Figure 11.**
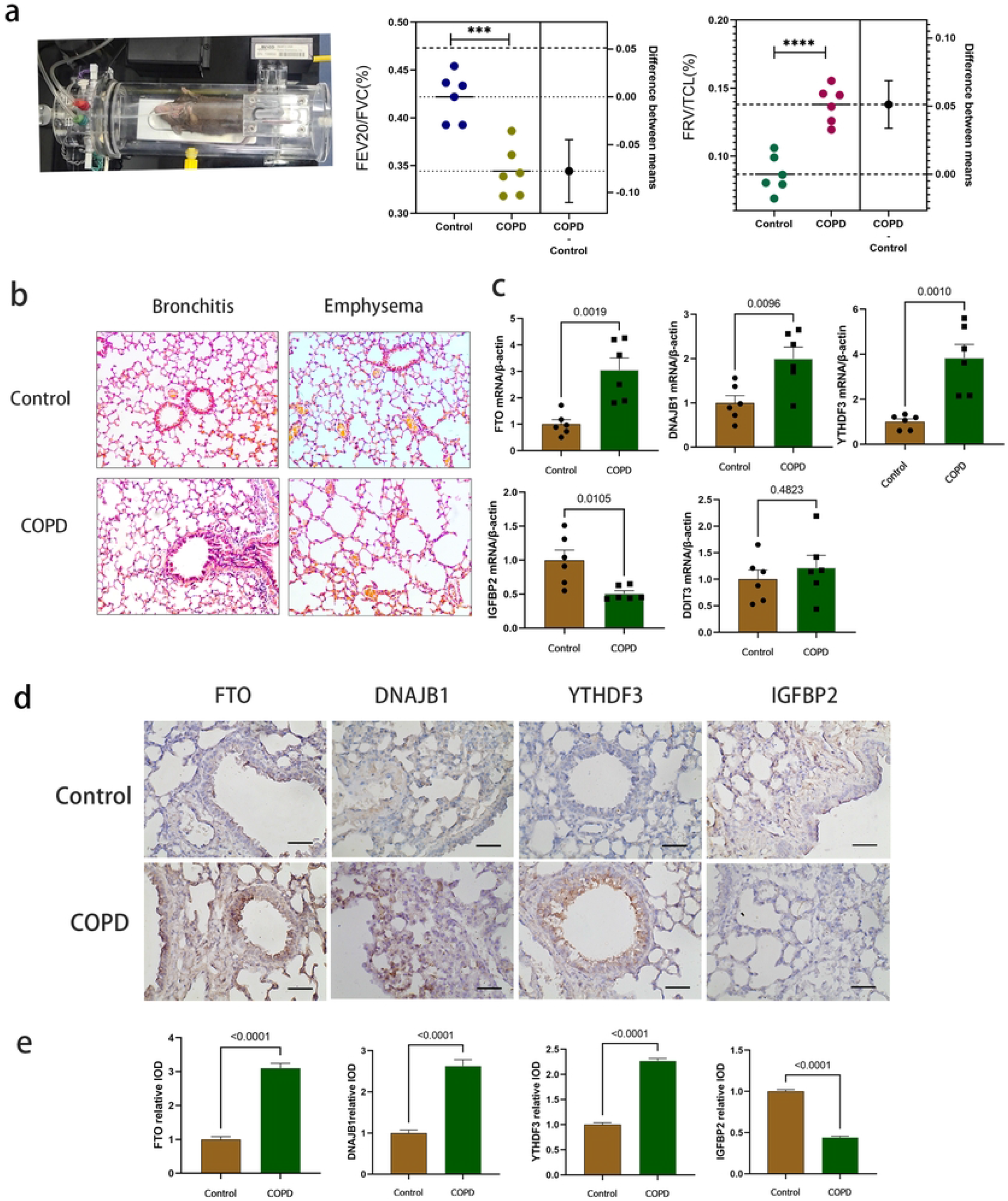
Experimental verification of the expression level of hub genes in mouse lung tissue. **(A):** COPD and control mice lung function detection. (B).COPD and control mice lung tissue HE staining (200×). **(C)**. The expression levels of five hub genes in lung tissues of COPD group and control group were verified by RT-qPCR (n =6/group). **(D)**.The expression levels of FTO, DNAJB1, YTHDF3 and IGF2BP2 in lung tissues of COPD group and control group were verified by immunohistochemistry. Scale bars represent 50 μm. **(E)** Immunohistochemical statistics: Compared with Control group P<0.0001. (***P<0.001 ****P<0.0001)

## 4. Discussion

Due to the global aging population and continued exposure to risk factors, the prevalence of chronic obstructive pulmonary disease (COPD) is expected to increase in the coming years (41) (42) (43). Recent studies have highlighted the involvement of RNA methylation and autophagy in the development and progression of COPD (9) (44). However, comprehensive comparative studies using high-throughput RNA sequencing data to investigate these processes in COPD are lacking. Identifying biomarkers associated with autophagy and methylation can aid in the diagnosis, management, and development of targeted therapies for COPD. This study investigated RNA methylation regulators using bulk RNA sequencing, identify hub genes using single-cell RNA sequencing (scRNA-seq), and validate the expression of key genes (*DDIT3, DNAJB1, FTO, IGF2BP2*, and *YTHDF3*) in COPD mouse models.

To study the association between RNA-modified genes and COPD in the GSE76705 dataset, 13 genes were identified through univariate logistic regression analysis. Among these genes, six were upregulated (*KIAA1429, IGF2BP2, TRMT61A, LRPPRC, HNRNPC, RRP8*), and eight were downregulated (*TRMT61B, TRMT10C, ZC3H13, METTL3, FMR1, YTHDF3, METTL*) in COPD. A co-expression network was constructed using weighted gene co-expression network analysis (WGCNA) to identify gene modules associated with RNA methylation and autophagy. Within the methylation pathway, the green and blue modules were screened and intersected with the 13 RNA-modified genes, leading to the identification of the YTHDF3 gene. YTHDF3 is a cytoplasmic m6A binding protein that has been implicated in the regulation of cytoplasmic metabolism of methylated mRNAs, particularly in the context of lung cancer (45). In cell experiments, knockdown of *YTHDF3* or *IGF2BP2* attenuated hypoxia/reoxygenation-induced injury in human bronchial epithelial cell by inactivating AKT, p38, NF-κB, and ERK1/2 pathways (46). The results of this study combined with our study suggest that *YTHDF3* gene is likely to play a promoting role in the occurrence and development of COPD. Interestingly, the expression levels of YTHDF3 showed a downregulation trend in an animal model of acute respiratory distress syndrome (ARDS) induced by LPS, which differs from the findings in the study on cigarette smoke-induced COPD (47). This inconsistent may be due to the difference in diseases and animal models.

Additionally, scRNA-seq analysis identified five highly infiltrating cell types (B-cells, CD4+ T-cells, CD8+ T-cells, macrophages, and monocytes) and differential gene analysis revealed the intersectant genes FTO and IGF2BP2 among the 21 clusters. FTO, the first identified RNA demethylase (48), has been associated with reduced lung function in recent genome-wide association studies (49). Animal experiments further showed upregulation of FTO in mice with LPS-induced ARDS(47). Similarly, emerging evidence has linked SNPs in the FTO gene to sarcopenia in COPD patients (50).

IGF2BP2, involved in post-transcriptional regulation, has been associated with type 2 diabetes and decreased glucose tolerance (51) (52). Knockdown of IGF2BP2 reduced injury in human bronchial epithelial cells and its binding to caspase 4 has been implicated in promoting airway inflammation and LPS-induced lung injury (46). However, experimental evidence for the role of YTHDF3 and IGF2BP2 in COPD is limited and further research is needed to elucidate their contributions to the disease.

GSEA and GSVA analyses revealed significant enrichment of differential genes in the autophagic pathway. Among the intersected autophagic markers, genes such as DNAJB9, NAMPT, DNAJB1, PPP1R15A, CDKN1A, CXCR4, DDIT3, VEGFA, FKBP1B, and NCKAP1 were identified. Increased expression of CXCR4 in CD27^−^ B-lymphocytes of smokers with COPD suggests its potential role in modulating neutrophil retention at inflammatory sites (53). This highlights CXCR4/CXCL12 signaling as a potential therapeutic target for evacuating neutrophils in chronic inflammatory diseases (54). The role of PPP1R15A in lung epithelial cells and lung mesenchymal cells has been explored. Specifically, loss of PPP1R15A has been observed to promote lung fibroblast senescence (55). These findings support the reliability and value of our data analysis. Autophagy plays a critical role in the lung’s inflammatory response, with proper regulation being essential for inhibiting pulmonary inflammation and facilitating leukocyte response to lung infections. However, dysregulated autophagy can lead to lung injury. Notably, the precise biological roles of NAMPT, DDIT3, DNAJB1, FKBP1B, and NCKAP1 genes in COPD remain to be determined, warranting further research.

GSEA revealed enrichment of immune cell-related pathways and autophagy-related signaling pathways in the differential analysis of the GSE76705 dataset. These pathways have been implicated in contributing to inflammatory damage in COPD. (56, 57). Additionally, functional enrichment analyses highlighted the significant connection of RNA methylation regulators to various regulatory processes, including translation, negative regulation of cellular macromolecules, and mRNA metabolism. These findings provide new insights into the mechanisms underlying COPD and offer a foundation for further research.

scRNA-seq profiles reveal heterogeneous inflammatory cells in COPD and help identify variable genes. Using SingleR, we identified 22 cell types, including upregulated B-cells, CD4+ T-cells, CD8+ T-cells, macrophages, and monocytes. T-cells and NK cells showed a negative correlation with macrophages and monocytes. Macrophages and monocytes had stronger interactions with B-cells, CD4+ T-cells, and CD8+ T-cells. We analyzed the output and incoming signals, finding that macrophages and monocytes play a significant role, particularly through IL-1, IL-6, and resistin pathways. Lung macrophages process tobacco smoke and particulate matter, leading to altered cytokine and chemokine release in COPD patients (58)(59). Autophagy scores indicated that enhanced macrophage RNA methylation and autophagy levels promote cellular communication, amplifying inflammation and contributing to COPD progression.

To verify the identified autophagy and methylated hub genes, we conducted animal experiments. Results showed significant upregulation of DNAJB1, FTO, and YTHDF3 mRNA expression, and downregulation of IGF2BP2 in COPD mouse lung tissue. Immunohistochemistry confirmed upregulation of FTO, YTHDF3, and DNAJB1 proteins, and downregulation of IGF2BP2 in COPD mouse lungs.

However, this study has limitations: small single-cell data sets, modest sample sizes, and the need for confirmation in future COPD patients.

In summary, RNA methylation and autophagy play a key role in COPD development, warranting further investigation into their mechanisms and the autophagy-mediated immune response in COPD pathogenesis.

## Author Contributions

LSX, SYX and OYY conceived and designed the study. LSX and SPP developed the methodology. ZLY, WYW, HSF, LMM, WDM, ZJ and LYT analyzed and interpreted the data. LSX, SPP, QW and WYW wrote, reviewed, and/or revised the manuscript. All authors contributed to the article and approved the submitted version.

## ACKNOWLEDGMENTS

We thank the authors who provided the GEO public datasets.

## Data Availability Statement

Data are available in a public, open access repository. Data are available on reasonable request. All data relevant to the study are included in the article or uploaded as supplemental information, The GEO datasets:

[GSE76705(https://www.ncbi.nlm.nih.gov/geo/query/acc.cgi?acc=GSE76705),

GSE167295(https://www.ncbi.nlm.nih.gov/geo/query/acc.cgi?acc=GSE167295),

GSE128033(https://www.ncbi.nlm.nih.gov/geo/query/acc.cgi?acc=GSE128033)]

